# Three water restriction schedules used in rodent behavioral tasks transiently impair growth and differentially evoke a stress hormone response without causing dehydration

**DOI:** 10.1101/2021.10.26.466027

**Authors:** Dmitrii Vasilev, Daniel Havel, Simone Liebscher, Silvia Slesiona-Kuenzel, Nikos K. Logothetis, Katja Schenke-Layland, Nelson K. Totah

## Abstract

Water restriction is commonly used to motivate rodents to perform behavioral tasks; however, its on hydration and stress hormone levels are unknown. Here, we report daily body weight and bi-weekly packed red blood cell volume and corticosterone in adult male rats across 80 days for three commonly used water restriction schedules. We also assessed renal adaptation to water restriction using post-mortem histological evaluation of renal medulla. A control group received ad libitum water. After one week of water restriction, rats on all restriction schedules resumed similar levels of growth relative to the control group. Nominal hydration was observed, and water restriction did not drive renal adaptation. An intermittent restriction schedule was associated with an increase in corticosterone relative to the control group. Our results suggest that the water restriction schedules used here will maintain welfare in rats. However, intermittent restriction evokes a stress response which could affect behavioral and neurobiological results. Our results also suggest that stable motivation in behavioral tasks may only be achieved after one week of restriction.

## Introduction

The use of water restriction to motivate rodents to perform goal-directed behavioral tasks has expanded in recent years due to the adoption of head-fixed behavioral paradigms using a water licking spout ^1–17^. Head-fixation has opened up new avenues of research in rodents, which were heretofore carried out largely in non-human primates, such as studies on visual perception ^18–23^, forelimb reaching ^7,16,24^, and arousal (using pupillometry) ^12,13,25,26^. Additionally, there has been a growing interest in using head-fixed rodents to study the neural representation of space using virtual reality ^27,28^. Head-fixation of rodents has also been used to gain access to membrane potentials during goal-directed behavior ^2,13,14^. Thus, the use of water restriction as a motivational tool for rodent behavioral paradigms will likely continue to be a fundamental tool in neuroscience.

However, the results of behavioral and neurobiological studies could be affected by the stress of water restriction, which has not been assessed in any of the water restriction schedules that are commonly used in neuroscience research. One study using rats limited access to water at 30 minutes per day and reported significant elevation of blood plasma ACTH and adrenal corticosterone (CORT) after 6 days ^29^. Training and measuring behavior and recording neuronal activity, on the other hand, can last many weeks of months and require water restriction well beyond a 6 days ^1,3,6,7,13,16,20,22^. Another study found no elevation of plasma CORT after 37 days during which rats were provided daily access to water for 15 minutes ^30^. These data suggest that the stress response may adapt during chronic water restriction at some point after 6 days. These studies have provided ‘snapshots’ of the stress response at 6 days and 37 days, but the change in CORT over the time course of a typical behavioral experiment remains unknown. Furthermore, these snapshots are limited to one type of restriction schedule, which limited water availability to unlimited volume consumption within a daily time window. To our knowledge, prior studies have not characterized the stress response to other types of schedules used in behavioral neuroscience, such as those that limit the total volume of water available per day. A clear picture of the stress response to the various water restriction schedules used in neuroscience research will enable the field to consider whether behavioral and neurobiological results might be affected by a stress response to water restriction.

The effect of water restriction schedules on hydration is also not well studied. The standard monitoring for dehydration in rodent behavioral neuroscience involves measuring reductions in body weight ^1,31– 33^. Importantly, body weight loss is not an ideal indicator of dehydration in rodents because their adaptive response to water scarcity is mild anorexia; by reducing food volume in the gastrointestinal tract, rodents reduce water lost through feces ^34–37^. Another standard assessment for dehydration is skin turgor ^32^. Although turgor is easy to deploy and offers a rapid clinical judgement of dehydration, it is subjective and only visible in stages of advanced dehydration. On the other hand, packed red blood cell volume (hematocrit, Hct) may be an objective clinical sign of dehydration ^38,39^. A prior study has shown that Hct in rats was elevated (indicative of dehydration) after 6 days of 30 minutes of daily water access ^29^. It remains unknown whether Hct is elevated during the chronic restriction that is used in behavioral tasks or whether Hct differs according to type of restriction schedule.

Here, we measured daily body weight and biweekly plasma CORT and Hct over 80 days in four groups of rats subjected to different water restriction schedules that are commonly used in behavioral studies ^1,3,6,7,13,16,20,22,32,40–46^. The restriction schedules were either ad libitum availability (control group), continuous volume-limited water, intermittent volume-limited water (i.e., alternating between 5 days of volume-limited daily water and 2 days of ad libitum access), or 30 minutes time-limited water. We found no evidence for dehydration or excessive stress response; however, the intermittent restriction schedule evoked a small stress response. We observed a 2-week adaptation period in which body weight is diminished in all three restriction groups and followed by normal growth. Kidney histology was used to measure changes in the renal medulla and demonstrated that these restriction schedules were not severe enough to drive long-term adaptation of the renal system. Overall, we found that months-long use of common restriction schedules in rats maintains rodent welfare, but that behavioral tests should avoid the initial 1 – 2 weeks of restriction during which motivation may be unstable and that intermittent schedules could potentially affect behavioral and neurobiological outcomes due to increased CORT.

## Results

We compared the effects three water restriction schedules on body weight, hematocrit (Hct), and blood plasma corticosterone (CORT) over 80 days. The restriction schedules were “timed”, “continuous”, and “intermittent.” The timed group was given 30 min of ad libitum access to water each day. The continuous group received approximately 12 mL per day as a single bolus. They began consuming this volume within seconds and, with a few drinking bouts interspersed with food consumption, the entire volume was consumed. If a rat in the continuous group lost more than 15% of their body weight in a week, then their daily water was increased by 2mL. Finally, the intermittent group received a repeating schedule of 12 mL of water per day for 5 days, followed by 48 hours of ad libitum water. On days with volume-limited water access, the consummatory behavior of these rats was noted as similar to that of the continuous group. The choice of volume and timing was based upon published literature and a detailed justification can be found in the methods section. A control group was monitored with ad libitum access to water for 80 days. Water administration occurred between 14:00 and 16:00. Prior to water administration, rats were weighed each day and blood was taken from the tail vein twice per week (usually Wednesday and Friday). Measurements were taken prior to water administration in order to capture the statuses of the rats in the water restricted state. Each group consisted of 6 male Sprague-Dawley rats. Rats were housed individually in order to control water intake. All rats were housed in the same room with cages randomly distributed across two racks of individually ventilated cages.

### Rats adapt to water restriction after two weeks and maintain normal hemotocrit levels

Body weight is frequently used as an indicator of overall health as well as an indirect measure of hydration status in rodents. We compared this measure across the three most frequently used restriction schedules. **Figure 1A** presents the average body weight in each group over 88 days. Water was removed on day 8. By day 88 (i.e., the 80^th^ day of water restriction), we observed significantly reduced body weights in all water restriction groups relative to the control group (**Figure 1B**). Body weight in the timed group was reduced by 16.2% relative to the ad libitum control group. The effect size and its 95% confidence intervals (ESCI) were at least an 8.5% decrease and at most a 22.5% decrease. Body weight loss in the continuous group was 28.4% (ESCI: between a 21.1% and a 34.3% loss). In the intermittent group, weight loss was 21.6% (ESCI: between a 12.7% and a 30.3% loss). A Bayesian ANOVA suggests that these data provide extremely strong evidence for a difference in weight between restriction schedules (BF = 3866.301). Post-hoc testing showed that rats on all water restriction schedules lost weight relative to the control group (BF’s for continuous, intermittent, and timed were 681.848, 23.841, and 15.394 respective to each group). The weight of rats on the continuous water restriction schedule was also lower than rats on the timed schedule (BF = 31.236). Water restriction clearly effected long-term body weight.

**Figure 1.**
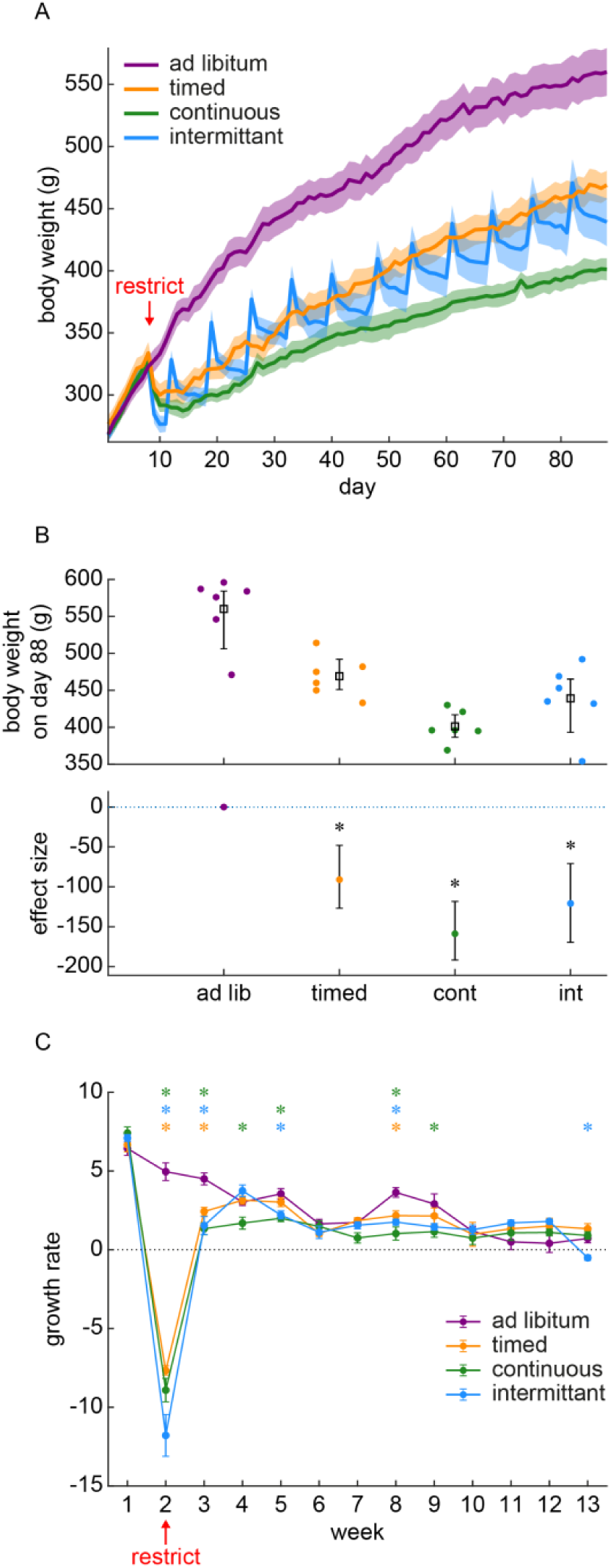
Water restriction evokes an overall body weight reduction that is due to weight loss during the first 2 weeks. **(A)** Body weight is plotted across the entire duration of the experiment (88 days). Water restriction began on day 8. The mean and standard error for each group of rats is shown as a line and shading. There were 6 rats per group. **(B)** The final body weight at the end of the experiment was reduced in all water restriction groups relative to the ad libitum control group. The upper panel shows the distribution of individual rats (dots) and the group mean and standard error. The lower panel shows the effect size relative to the ad libitum control group and the error bars show the 95% confidence interval for the effect size. **(C)** Weekly body weight change is plotted throughout the experiment. The data markers and lines show the mean and standard error for each group of rats. Negative growth (below the dotted line) indicates weight loss, whereas positive points indicate growth. Post-hoc Bayesian t-tests on the alternative hypothesis that control group growth was greater than the restricted group’s growth is illustrated with a star when BF is greater than 3. The color of the star indicates the identity of the group being compared against the ad libitum control group. A BF>3 indicates that these data provide evidence supporting the alternative hypothesis that growth in the control group was greater than that of the restricted group. The growth differences occurred primarily during the first 2 weeks of restriction.

Long-term weight loss may indicate a long-term disruption of rodent health. However, in **Figure 1A**, it appears that much of the weight loss occurs during the first two weeks and that growth normalizes thereafter. It is, therefore, possible that the lower weights after 80 days of restriction were due to a brief period of weight loss occurring at the start of water restriction and that, although these early losses were never re-gained, growth proceeded normally. We formally examined this question by measuring growth as weekly body weight change (**Figure 1C**). We found that growth largely normalized after 2 weeks of water restriction. There was an interaction between restriction schedule and week (Bayesian repeated measures ANOVA: BF = 3.198×10^118^), which was due to body weight losses that occurred largely in weeks 2 and 3. Thus, our results suggest that the large decrease in body weight after 80 days of chronic water restriction is not due to long-term growth impairment. Instead, since growth normalized after two weeks of restriction, it is likely that this brief window of weight loss is followed by adaptation to the new environmental demands. It is possible however that, prior to adapting, the weight loss during the first two weeks is due to dehydration.

We used hematocrit (Hct) levels to more directly assess whether the first two weeks of water restriction were associated with a change in hydration. Hct was measured as the percent packed cell volume in centrifuged blood samples taken twice per week (**Figure 2A**). Hct differed over time (Bayesian ANOVA interaction between time and schedule: BF = 1.944×10^55^). Hct was increased during the first week of restriction, which was also the first week in which a blood sample was obtained. However, this increase occurred in the control group which suggests that this change was not specific to water restriction. **Figure 2B** shows the Hct values recorded across 80 days of chronic water restriction. The data provide strong evidence supporting the null hypothesis that mean Hct did not differ between restriction schedules (BF = 0.052). Therefore, the drop in body weight during the first two weeks of water restriction is not due to a change in hydration. Instead, the reason for the weight loss during the initial two weeks of water restriction may be a mild anorexic response that reduces water loss through feces. As part of this adaptive response, the renal mechanisms for water conservation may be engaged so that rats can resume normal growth (despite limited water availability) beginning in the third week of restriction (see **Figure 1C**).

**Figure 2.**
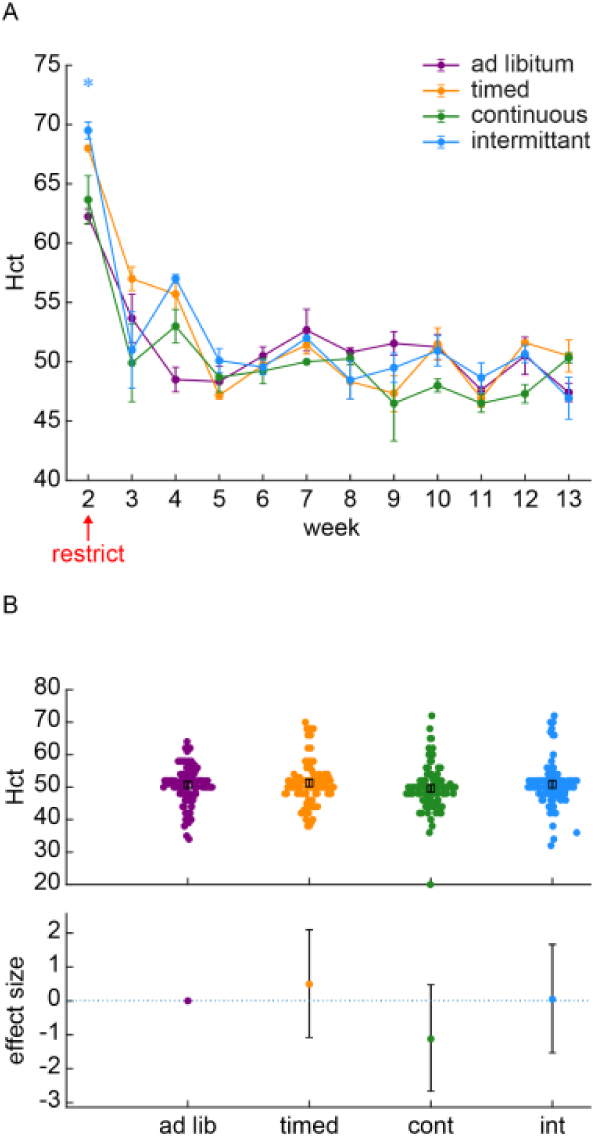
Hct did not differ between the ad libitum control group and groups of rats subjected to various water restriction schedules. **(A)** % Hct is plotted as the group average of all samples collected each week. The error bars indicate the standard error. Although there in an increase in Hct during the first week of blood collection, this occurred in all groups inclusive of the control group. **(B)** An assessment of effect sizes comparing all Hct values collected over 88 days suggests that Hct does not differ between groups. The effect sizes (95% confidence interval) relative to the ad libitum control group were: timed group – 1.0% (−2.1 to +4.1%), continuous group – 2.2% (−5.2 to +0.9%), intermittent group – 0.1% (−3.0 to +3.3%).

### Water restriction evokes a stress response in rats on an intermittent water restriction schedule

Adaptation to water restriction through an anorexic response may increase circulating stress hormones, given that food restriction can elevate corticosterone (CORT) in rodents ^47,48^. We measured blood plasma CORT twice per week (**Figure 3A**). CORT values are reported starting from week 3 because inadequate blood volumes were obtained in week 1 (prior to restriction) and week 2 (start of restriction). There was an interaction between restriction schedule and time (BF = 1.353×10^13^), which were driven by an early increase in the intermittent group (post-hoc Bayesian t-tests, BF>3, except BF = 2.082 for week 5 and BF = 2.423 for week 9). Given that week 3 was associated with reduced growth (see **Figure 1C**) and potential anorexia in all restriction groups, the specificity of the stress response for the intermittent restriction group suggests that an initial anorexic response during the first 2 weeks of water restriction is not associated with a stress response.

**Figure 3.**
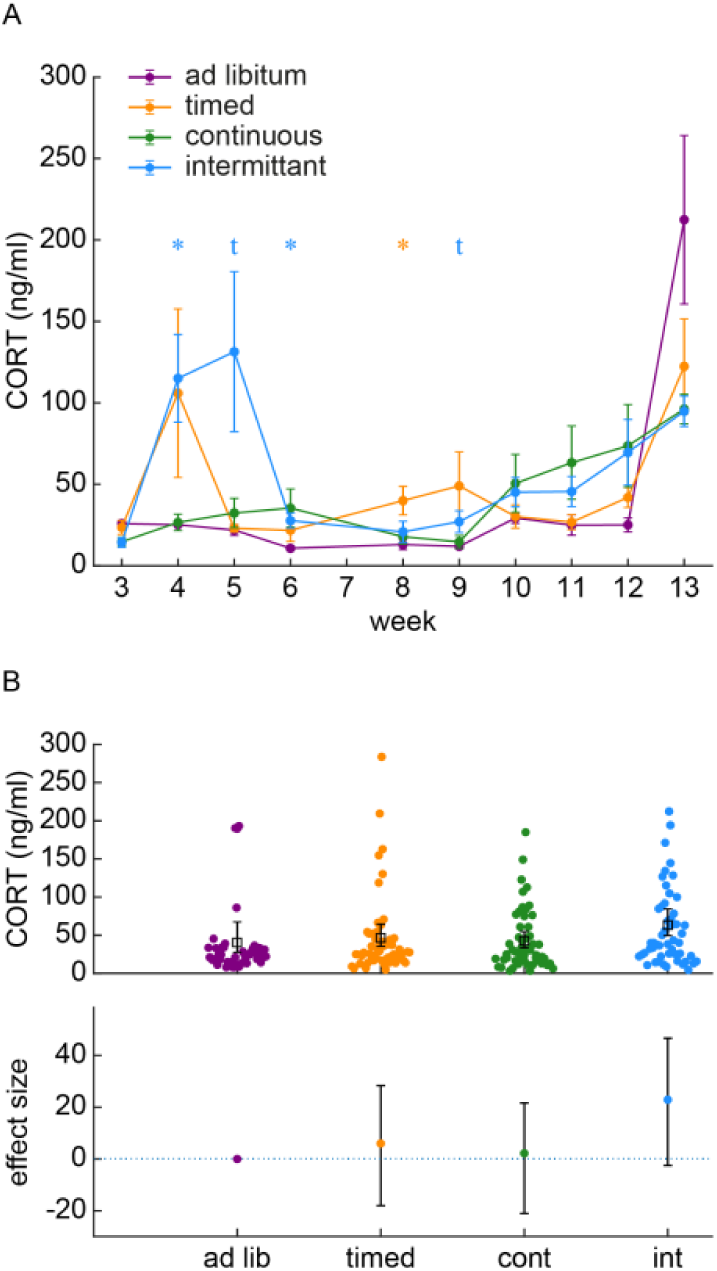
CORT is increased during intermittant water restriction. **(A)** The average and standard error of weekly CORT values are plotted for each group of rats. The intermittant group has occasionally elevations in CORT. **(B)** Collapsing all measurements over time revealed that the intermittant group has elevated CORT relattive to the ad libitum control group. The upper panel shows individual data points and the lower panel shows the effect sizes of group differences relative to the ad libitum control group.

On the other hand, a stress response may also be evoked by environmental instability which occurs specifically in the intermittent group. Environmental instability occurs in the intermittent group because these rats repeatedly encountered water losses after the periodic 2-day breaks from restriction. The instability in the environment altered the body state of these rats by producing a highly variable ‘sawtooth’ pattern in intermittent group body weights (see **Figure 1A**). Our data suggest that the environmental instability encounted by the intermittent schedule group could be a psychological stressor that evokes a chronic stress response. We assessed this by collapsing CORT measurements from all timepoints to assess whether CORT was overall higher in the intermittent group (**Figure 3B**). The ESCI in the timed group spanned from a 46% reduction to a 69.4% elevation in CORT, which suggests no effect of timed restriction on CORT. Similarly, the continuous group ESCI ranged from a 51.8% reduction in CORT up to a 53.0% increase in CORT. However, the intermittent group was associated with an effect size of 56.6% with the ESCI ranging from roughly no change (−6.9%) up to a 115.0% increase. CORT in the ad libitum control group was 40±9 ng/mL, whereas in the intermittent group it increased to 63±9 ng/mL. Given that the ESCI is the 95% confidence interval for the effect size, the average CORT was likely increased specifically in the intermittent group. We formally tested the alternative hypothesis that the intermittent group had higher CORT than the ad libitum control group using a Bayesian Mann-Whitney U Test (due to the skew of the intermittent group distribution). A BF of 27.47 suggested that the collected data may be taken as strong evidence in favor of the alternative hypothesis. Therefore, the intermittent water restriction schedule used here may present a psychological stressor that evokes a significant increase in CORT that is around 56.6% higher than under ad libitum conditions.

### Water restriction is not associated with adaptation of the Loops of Henle

In response to water scarcity, organisms adapt by producing hyperosmotic urine. This physiological adaptation depends upon the lengths of the Loops of Henle in the renal medulla ^49^. There is evidence that structural adaptation of the Loops of Henle can occur over the timescale of a few weeks during water deprivation ^50^. We assessed whether the water restriction schedules used here were severe enough to promote structural changes to the kidneys using post-mortem histological measurements of relative medullary thickness (RMT). RMT is indicative of a shift in the size of the renal medulla relative to the cortex indicating lengthening of the Loops of Henle ^51^. An increased RMT is associated with urine osmolality and can therefore be used as a surrogate measure of an organism’s ability to produce hyperosmotic urine in response to water scarcity ^49,52,53^. Kidney sections were inspected, and measurements made by an individual blind to the water restriction group assignments of the rats. RMT was measured according to two formulae that capture changes in the size of the renal medulla relative to the rest of the kidney (**Figure 4A**). We found that neither of these measures differed across groups (**Figure 4B, 4C**). A Bayesian one-way ANOVA suggested moderate support for the null hypothesis for similar outer medulla to cortex ratio (OMR) across groups (BF = 0.23). The result of the Bayesian one-way ANOVA for total medulla to cortex ratio (MR) data was ambiguous, but close to the threshold for moderate evidence supporting the null hypothesis (BF = 0.66). Taken together, it is unlikely that water restriction schedules evoked lengthening of the Loops of Henle. Interestingly, there was a significant difference in post-mortem kidney weight across groups (Bayesian one-way ANOVA, BF = 7.514). The data suggest that, in the continuously restricted group, kidney weight was reduced (**Figure 4D**). The continuous group kidney weight was 16.1% lower than the kidneys in the ad libitum control group with a 95% confidence interval of an 8.5% loss up to a 23.0% loss. Post-hoc Bayesian t-tests indicated that the kidney weights of rats in the continuous water restriction group were significantly lower than those of the ad libitum group (BF = 13.08), as well as the intermittent water restriction group (BF = 14.35) and the timed group (BF = 5.377). Collectively, these data suggest that water restriction is not severe enough to evoke structural adaptations of the renal medulla; however, continuous water restriction may lead to a modification of gross kidney mass.

**Figure 4.**
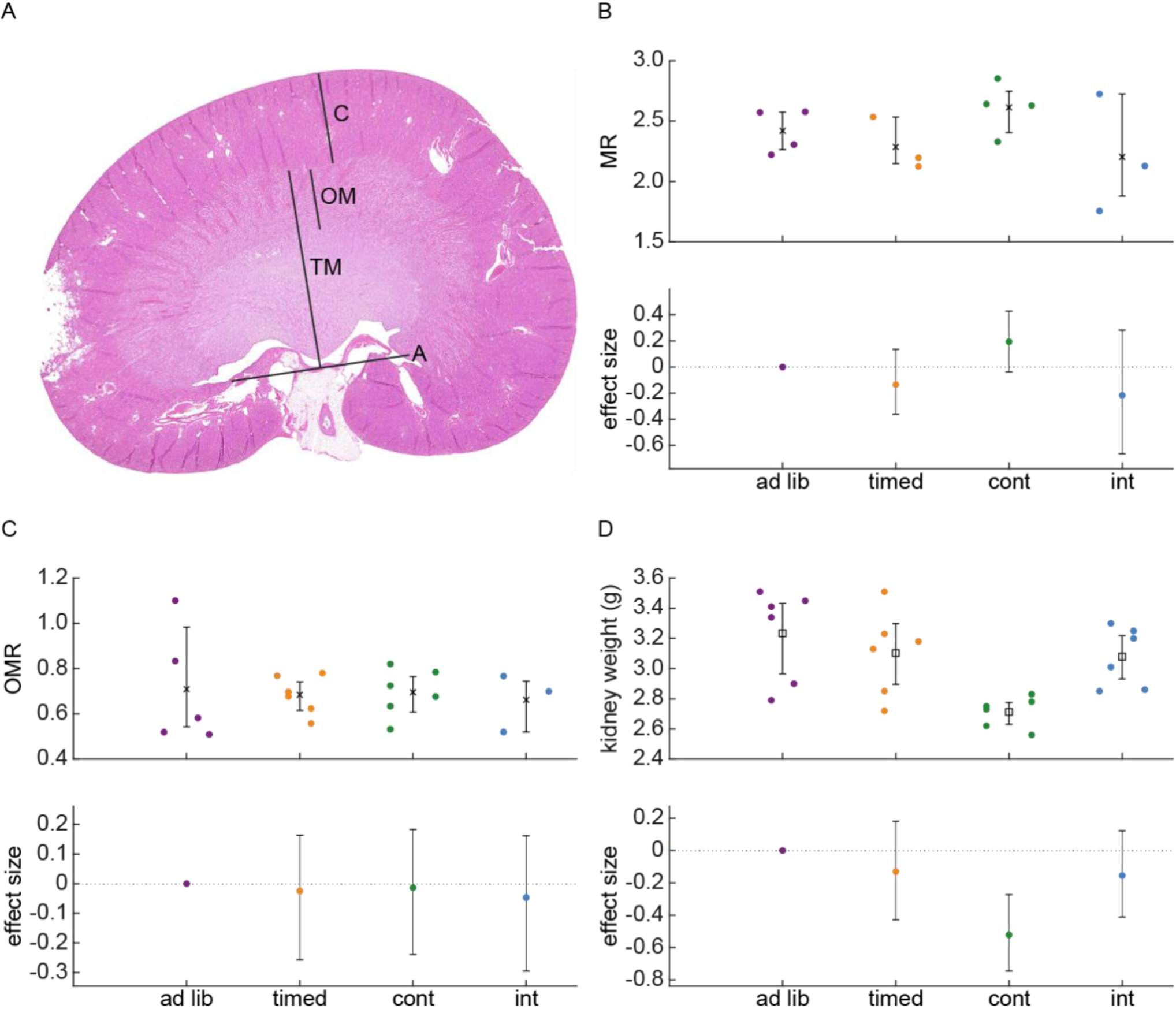
Water restriction is not associated with alterations in the Loops of Henle. **(A)** This example tissue section of the kidney shows the gross anatomical makers used to delineate the cortex (C), the outer medulla (OM), the total medulla (TM) demarcated as the distance between the capsule and the ‘assisting line’ (A). The assisting line connects the two points where the ureter connects to the kidney and was used to have a standardized starting point for measurements. The TM may or may not include the entire inner medulla, which sometimes crossed the assisting line. We assessed relative medullary thickness using two formulae. The first was the OM to C ratio (OMR) and the other was the TM to C ratio (MR). **(B, C)** Neither the MR (B) nor the OMR (C) differed across groups of rats. The plots show the individual data points where complete sections could be obtained to make a clear assessment of these measures. The lower panel shows the effect sizes relative to the ad libitum control group and the 95% confidence intervals of those effect sizes. **(D)** Post-mortem kidney weight was measured and indicated a reduction in kidney weight in the group of rats subjected to continuous water restriction. The upper panel shows individual data points and the lower panel shows effect sizes and confidence intervals relative to the ad libitum control group.

## Discussion

Water restriction is a widely used tool in neuroscience research in rodents; however, effects of these water restriction schedules on objective measures of hydration and on stress hormone level are unknown. It is also possible that rodents readily adapt to water restriction by structural modification of the kidneys. Here, we measured daily body weight and biweekly plasma corticosterone (CORT) and packed red blood cell volume (hematocrit, Hct) over 80 days in four groups of rats subjected to different water restriction schedules that are commonly used in behavioral studies. We found no evidence for changes in Hct. Although stress hormones were also not generally altered by water restriction, the intermittent group had a minor elevation in CORT that may be a behaviorally and neurobiologically-relevant stress response. We also observed a 1 – 2 week period in which body weight is diminished in all three restriction groups and followed by normal growth, which could result in unstable baseline motivation for water reward during this adaptive period. Kidney histology was used to measure changes in the renal medulla and demonstrated that these commonly used restriction schedules are not severe enough to drive long-term adaptation of the renal system. However, we cannot exclude the possibility that renal aquaporin expression may have adapted in response to water restriction, given that plasticity of aquaporin expression has been demonstrated in the response of rodents to changes in seasonal water scarcity in the wild ^54^. Overall, we found that months-long use of common restriction schedules in rats maintains rodent welfare. Our results suggest that behavioral tests should avoid the initial 1 – 2 weeks of adaptation and that intermittent schedules may evoke a stress response that affects behavioral and neurobiological outcomes.

### Implications for rodent welfare

The welfare of rodents is a key objective in all experiments primarily for ethical reasons, but also because unhealthy animals cannot yield normal data. Water restriction could affect welfare by evoking dehydration; however, our results suggest that the restriction schedules used here do not cause dehydration. We assessed hydration by measuring Hct. Typical Hct values in adult, male rats have been reported to range between 33 and 57 ^55^ and around 42 in rats that are not water restricted ^56^. Although Hct depends on age and body weight, it largely stabilizes between 40 to 45 in rats of the age and body weight used in the present study ^57^. Therefore, we observed normal Hct values that are typical for non-water restricted rats of this gender and age. Our findings suggest that hydration is not affected by any of the water restriction scheduled used in the present study. Normal hydration is presumably maintained by the rats via renal adaptation and the production of hyperosmotic urine.

### Implications for maintaining stable motivation of rodents during goal-directed behavioral tasks

The schedules tested here are commonly used in rodent behavioral neuroscience experiments. For example, some laboratories have chosen to use continuous volume-limited restriction because motivation is reduced after each break in an intermittent schedule ^32,58,59^. However, intermittent schedules are also common ^1,32,60^. Motivation can also be maintained at a stable level with a timed access schedule. Various laboratories have motivated behavior by time-limited water access from 10 min per day to 1 hour per day ^3,17,40–42^. Our finding that body growth is temporarily reduced for the initial ∼2 weeks of water restriction for all schedule types suggests that this is a period of adaptation. During this period when rodents adapt to the new environmental constraints, they may have increased motivation to collect reward. Our results suggest that collection of behavioral and neurobiological data during this period may include instabilities in how rodents respond to water rewards. Therefore, allowing this adaptation period to pass before collecting data may be a useful practice in behavioral neuroscience research.

Other recent methods of water restriction that provide less palletable 2% citric acid water ad libitum in the home cage have not been assessed for stress hormone release or kidney adaptation ^31,33^. However, it is likely that our findings would be similar under those conditions because rodents given citric acid in water effectively self-restrict their water intake (due to the aversive taste of the water) to approximately the same volume of daily water provided in our study to the continuous restriction and intermittent restriction groups ^33^. Although providing ad libitum citric acid water is less labor intensive and is an efficient way to motivate rodents without needing to administer precise daily allotments of water tailored to individual rats, the use of citric acid results in fewer daily trials performed compared to water restriction ^33^. Therefore, the water restriction schedules used here may be most relevant to studies that aim to maximize the number of trials performed.

### Implications of the stress response in rats on an intermittent water restriction schedule

Intermittent water restriction could be a physiological stressor due to the saw-tooth pattern of repeated weight loss and weight rebound. It could also be a psychological stressor due to unstable environmental water availability. We observed a mean increase of plasma CORT to 63 ng/mL in rats on the intermittent restriction schedule. The blood plasma collection procedure (single tail vein puncture) did not affect the measurements because CORT requires ∼20 min to elevate after a tail puncture ^61^. Our data support the notion that intermittent water restriction is a stressor, but it is a relatively minor stressor compared to air puff startle (450 ng/mL ^62^), restraint stress (250 to 800 ng/mL ^63,64^) and forced swimming (400 to 500 ng/mL ^65^). However, CORT level during an intermittent water restriction schedule is similar to the CORT increase after handling in rats subjected to early life maternal separation stress (100 ng/mL ^66^) and the stress response to environmental noise (100 to 200 ng/mL ^65^). In sum, our results demonostrate that the water restriction schedules used in this study provided appropriate levels of hydration, but that rats on an intermittent water restriction schedule have a stress response which may have behavioral, neurochemical, and neurophysiological effects.

## Methods

### Subjects

Experiments were carried out with 24 male Sprague-Dawley rats (specific pathogen free, Charles River Laboratories, Sulzfeld, Germany). Rodents were single housed to control water administration. An 08:00 to 20:00 lights on cycle was used so that data could be carried out under normal lighting for the researchers. All experiments were carried out with approval from the local authorities and in compliance with the German Law for the Protection of Animals in experimental research and the European Community Guidelines for the Care and Use of Laboratory Animals.

### Water restriction procedures

Rats were divided into 4 groups covering 3 different water restriction schedules and an ad libitum control group. The restriction schedules were “timed”, “continuous”, and “intermittent.” The timed group was given 30 min of ad libitum access to water each day. The continuous group received approximately 12 mL per day. This small volume of water was delivered using a custom-made water bottle that released water only during consumption. If a rat in the continuous group lost more than 15% of their body weight in a week, then their daily water was increased by 2mL. The intermittent group received a repeating schedule of 12 mL of water per day for 5 days, followed by 2 days of ad libitum water.

The 12 mL volume of water for the continuous and intermittent groups was chosen based on our experience motivating behavioral task performance by head-fixed rats, as well as the physiological needs of adult male rats. Typical water ingestion patterns of the adult (300 – 400 g) male rat consist of consuming 20 – 30 mL per day when it is freely (ad libitum) available ^1,67^. However, rodents have highly effective renal mechanisms for water conservation, which allow them to remain hydrated and healthy when consuming less than 20 – 30 mL per day. For example, under conditions in which rats were allowed to consume as much water as they would like in their home cage, but requiring them to perform physical effort for access by pressing a lever, their daily water consumption was lower (15 mL) per day ^68^. This daily amount (15 mL consumed by 300 g rats) is approximately equivalent to the requirement to prevent cellular dehydration derived from fluid maintenance formulas, which calculate a requirement of 50 mL / kg of body weight per day to maintain normal hydration ^67^. Further reductions below ∼ 15 mL per day for a 300 g rat will activate renal mechanisms allowing rats to conserve water and remain hydrated; therefore, 50 mL/kg/day reflects an upper limit to the amount of water that must be allocated on a water restriction schedule. Water restriction protocols are generally designed to reduce water availability below this upper limit because increased water restriction is associated with higher goal-directed behavioral task performance as measured by percent correct choices in a sensory stimulus discrimination task ^32^. In addition to the observation of this phenomenon by Guo et al., we have also observed in our own unpublished head-fixed rat behavioral experiments that rats are not motivated unless they receive 60 to 80% of this upper limit (i.e., 9 to 12 mL total water per day). For example, we have observed that rats who receive 14 - 17mL will, in the next behavior session, omit (not perform) a large proportion of trials (6 to 33%). Therefore, providing too much water will reduce their motivation and they will not perform the task. Thus, we have found that 10 mL is adequate for most animals to be motivated to perform the task, but some animals must receive only 8 mL. It is possible that rats who require only 8mL of water per day to perform the task may have stronger physiological mechanisms for water conservation. In general, rats consume 3 – 8 mL during the behavioral task and the remaining amount (up to 8 - 12 mL) is provided in the cage. Therefore, for the present experiments, we chose to test 12 mL of daily water allotment.

### Method of blood sampling and measurement of CORT and Hct

A small blood sample (∼ 0.25 mL) was taken from the tail vein without anesthesia while the rat was held in a restraint tube. Samples were collected bi-weekly. During 2 weeks prior to starting water restriction, the animals were handled and habituated to restraint to reduce the stress response during data collection. Blood samples for Hct measurement were collected in a capillary tube and immediately centrifuged. The packed cell volume was measured against a chart calibrated for the capillary. Blood was centrifuged and the blood plasma was harvested and stored at −80 C. Blood plasma CORT was measured by a commercial firm using ELISA kits (Idexx Laboratories, Ludwigsburg, Germany).

### Kidney histology

Rat kidneys were freshly fixed in 4% PFA, washed in RNase free water and transferred in 70% RNase free ethanol. Kidneys were bisected longitudinal before automated embedding in paraffin using a STP120 (Thermo Fisher Scientific). Each paraffin-embedded half was sectioned (10µm sections) using a microtome HM340E (Thermo Fisher Scientific).

Histological staining was performed on deparaffinized and hydrated serial sections of rat kidney. Hematoxylin and Eosin (H&E) staining visualized cell nuclei (black, dark blue) and counterstains cytoplasm and connective tissue fibers (different shades of pink). In detail (also shown in Table 1), staining was started by deparaffinization with two steps of absolute Xylene followed by rehydration steps with a descending ethanol row. Hematoxylin staining was done with Mayer’s hematoxylin solution (Carl Roth GmbH, T865.1) for 10 minutes followed by 10 minutes bluing in lukewarm running tap water. Counterstain was done with 1% Eosin Y solution (Carl Roth GmbH, 3137.2) for 2 minutes followed by a differentiation step in 70% Ethanol for 30 seconds. Stained sections were mounted with Roti-Histokitt (Carl Roth GmbH, 6638.1). Stained sections were stored at room temperature until imaging analysis was performed.

**Table 1:** Deparaffinization, Rehydration and H&E staining procedure

|  |  |  |
| --- | --- | --- |
| **10 min** | Xylene I | Deparaffinization |
| **10 min** | Xylene II |  |
| **5 min** | Ethanol absolute I | Rehydration |
| **5 min** | Ethanol absolute II |  |
| **5 min** | Ethanol 96% I |  |
| **5 min** | Ethanol 96% II |  |
| **5 min** | Ethanol 70% I |  |
| **5 min** | Ethanol 70% II |  |
| **5 min** | Ethanol 50% I |  |
| **5 min** | Ethanol 50% II |  |
| **5 min** | distilled water |  |
| **10 min** | Mayer's hematoxylin | Nuclear staining |
| **30 s** | distilled water |  |
| **10 min** | running lukewarm tap water | Bluing |
| **30 s** | distilled water |  |
| **2 min** | Eosin Y 1% | Counterstaining |
| **30 s** | distilled water |  |
| **30 s** | Ethanol 70% | Dehydration |
| **3 min** | Ethanol 96% |  |
| **3 min** | Ethanol absolute I |  |
| **3 min** | Ethanol absolute II |  |
| **3 min** | Roti-Histol I | Clearance before mounting |
| **3 min** | Roti-Histol II |  |
|  | Roti-Histokitt | Mounting |

## References

1. Schwarz, C. et al. The head-fixed behaving rat--procedures and pitfalls. Somatosens Mot Res 27, 131 148 (2010).

2. Sachidhanandam, S., Sreenivasan, V., Kyriakatos, A., Kremer, Y. & Petersen, C. C. H. Membrane potential correlates of sensory perception in mouse barrel cortex. Nat Neurosci 16, 1671 1677 (2013).

3. Scott, B. B., Brody, C. D. & Tank, D. W. Cellular Resolution Functional Imaging in Behaving Rats Using Voluntary Head Restraint. Neuron 80, 371 384 (2013).

4. Lee, S.-H. et al. Activation of specific interneurons improves V1 feature selectivity and visual perception. Nature 488, 379 383 (2012).

5. Jurjut, O., Georgieva, P., Busse, L. & Katzner, S. Learning Enhances Sensory Processing in Mouse V1 before Improving Behavior. J Neurosci 37, 6460 6474 (2017).

6. Runyan, C. A., Piasini, E., Panzeri, S. & Harvey, C. D. Distinct timescales of population coding across cortex. Nature 548, 92 96 (2017).

7. Yttri, E. A. & Dudman, J. T. Opponent and bidirectional control of movement velocity in the basal ganglia. Nature 533, 402 406 (2016).

8. Stringer, C. et al. Spontaneous behaviors drive multidimensional, brainwide activity. Science 364, eaav7893 (2019).

9. Schröder, S. et al. Arousal Modulates Retinal Output. Neuron 107, 487-495.e9 (2020).

10. Steinmetz, N. A., Zatka-Haas, P., Carandini, M. & Harris, K. D. Distributed coding of choice, action and engagement across the mouse brain. Nature 576, 266–273 (2019).

11. Peters, A. J., Fabre, J. M. J., Steinmetz, N. A., Harris, K. D. & Carandini, M. Striatal activity topographically reflects cortical activity. Nature 591, 420–425 (2021).

12. Reimer, J. et al. Pupil fluctuations track fast switching of cortical states during quiet wakefulness. Neuron 84, 355 362 (2014).

13. McGinley, M. J., David, S. V. & McCormick, D. A. Cortical Membrane Potential Signature of Optimal States for Sensory Signal Detection. Neuron 87, 179 192 (2015).

14. Polack, P.-O., Friedman, J. & Golshani, P. Cellular mechanisms of brain state-dependent gain modulation in visual cortex. Nat Neurosci 16, 1331 1339 (2013).

15. Zhang, S. et al. Long-range and local circuits for top-down modulation of visual cortex processing. Science 345, 660–665 (2014).

16. Mathis, M. W., Mathis, A. & Uchida, N. Somatosensory Cortex Plays an Essential Role in Forelimb Motor Adaptation in Mice. Neuron 93, 1493-1503.e6 (2017).

17. Foo, C. et al. Reinforcement learning links spontaneous cortical dopamine impulses to reward. Curr Biol (2021) doi:10.1016/j.cub.2021.06.069.

18. Sriram, B., Meier, P. M. & Reinagel, P. Temporal and spatial tuning of dorsal lateral geniculate nucleus neurons in unanesthetized rats. J Neurophysiol 115, 2658–2671 (2016).

19. Khastkhodaei, Z., Jurjut, O., Katzner, S. & Busse, L. Mice Can Use Second-Order, Contrast-Modulated Stimuli to Guide Visual Perception. J Neurosci 36, 4457 4469 (2016).

20. Jin, M. & Glickfeld, L. L. Mouse Higher Visual Areas Provide Both Distributed and Specialized Contributions to Visually Guided Behaviors. Curr Biol 30, 4682-4692.e7 (2020).

21. Jacobs, E. A. K., Steinmetz, N. A., Peters, A. J., Carandini, M. & Harris, K. D. Cortical State Fluctuations during Sensory Decision Making. Curr Biol 30, 4944-4955.e7 (2020).

22. Laboratory, T. I. B. et al. Standardized and reproducible measurement of decision-making in mice. Elife 10, e63711 (2021).

23. Pakan, J. M. P., Currie, S. P., Fischer, L. & Rochefort, N. L. The Impact of Visual Cues, Reward, and Motor Feedback on the Representation of Behaviorally Relevant Spatial Locations in Primary Visual Cortex. Cell Reports 24, 2521–2528 (2018).

24. Galiñanes, G. L., Bonardi, C. & Huber, D. Directional Reaching for Water as a Cortex-Dependent Behavioral Framework for Mice. Cell Reports 22, 2767–2783 (2018).

25. Reimer, J. et al. Pupil fluctuations track rapid changes in adrenergic and cholinergic activity in cortex. Nat Commun 7, 13289 (2016).

26. Vinck, M., Batista-Brito, R., Knoblich, U. & Cardin, J. A. Arousal and locomotion make distinct contributions to cortical activity patterns and visual encoding. Neuron 86, 740 754 (2015).

27. Harvey, C. D., Collman, F., Dombeck, D. A. & Tank, D. W. Intracellular dynamics of hippocampal place cells during virtual navigation. Nature 461, 941–946 (2009).

28. Radvansky, B. A. & Dombeck, D. A. An olfactory virtual reality system for mice. Nat Commun 9, 839 (2018).

29. Arnhold, M. M., Yoder, J. M. & Engeland, W. C. Subdiaphragmatic Vagotomy Prevents Drinking-Induced Reduction in Plasma Corticosterone in Water-Restricted Rats. Endocrinology 150, 2300–2307 (2009).

30. Heiderstadt, K. M., McLaughlin, R. M., Wright, D. C., Walker, S. E. & Gomez-Sanchez, C. E. The effect of chronic food and water restriction on open-field behaviour and serum corticosterone levels in rats. Laboratory animals 34, 20 28 (2000).

31. Urai, A. E. et al. Citric Acid Water as an Alternative to Water Restriction for High-Yield Mouse Behavior. Eneuro 8, ENEURO.0230-20.2020 (2021).

32. Guo, Z. V. et al. Procedures for behavioral experiments in head-fixed mice. Plos One 9, e88678 (2014).

33. Reinagel, P. Training Rats Using Water Rewards Without Water Restriction. Front Behav Neurosci 12, 84 (2018).

34. Desai, M., Gayle, D., Kallichanda, N. & Ross, M. G. Gender specificity of programmed plasma hypertonicity and hemoconcentration in adult offspring of water-restricted rat dams. J Soc Gynecol Invest 12, 409 415 (2005).

35. Rowland, N. E. Food or fluid restriction in common laboratory animals: balancing welfare considerations with scientific inquiry. Comparative medicine 57, 149 160 (2007).

36. Bekkevold, C. M., Robertson, K. L., Reinhard, M. K., Battles, A. H. & Rowland, N. E. Dehydration parameters and standards for laboratory mice. J Am Assoc Laboratory Animal Sci Jaalas 52, 233–9 (2013).

37. Watts, A. G. Dehydration-Associated Anorexia Development and Rapid Reversal. Physiol Behav 65, 871–878 (1999).

38. Dorrington, KeithL. SKIN TURGOR: DO WE UNDERSTAND THE CLINICAL SIGNã Lancet 317, 264–266 (1981).

39. Hansen, B. & DeFrancesco, T. Relationship between hydration estimate and body weight change after fluid therapy in critically ill dogs and cats. J Vet Emerg Crit Car 12, 235–243 (2002).

40. Barnet, R. C., Cole, R. P. & Miller, R. R. Temporal integration in second-order conditioning and sensory preconditioning. Anim Learn Behav 25, 221–233 (1997).

41. Denniston, J. C., Cole, R. P. & Miller, R. R. The role of temporal relationships in the transfer of conditioned inhibition. Journal of experimental psychology. Animal behavior processes 24, 200 214 (1998).

42. Laraway, S., Snycerski, S. & Poling, A. MDMA and Performance Under a Progressive-Ratio Schedule of Water Delivery. Exp Clin Psychopharm 11, 309 (2003).

43. Panigrahi, B. et al. Dopamine Is Required for the Neural Representation and Control of Movement Vigor. Cell 162, 1418 1430 (2015).

44. Jaramillo, S. & Zador, A. M. The auditory cortex mediates the perceptual effects of acoustic temporal expectation. Nat Neurosci 14, 246 251 (2011).

45. Fujisawa, S., Amarasingham, A., Harrison, M. T. & Buzsáki, G. Behavior-dependent short-term assembly dynamics in the medial prefrontal cortex. Nat Neurosci 11, 823 833 (2008).

46. Pan, W.-X., Brown, J. & Dudman, J. T. Neural signals of extinction in the inhibitory microcircuit of the ventral midbrain. Nat Neurosci 16, 71 78 (2012).

47. Heiderstadt, K. M., McLaughlin, R. M., Wrighe, D. C., Walker, S. E. & Gomez-Sanchez, C. E. The effect of chronic food and water restriction on open-field behaviour and serum corticosterone levels in rats. Lab Anim 34, 20–28 (2000).

48. Abrahamsen, G. C., Berman, Y. & Carr, K. D. Curve-shift analysis of self-stimulation in food-restricted rats: relationship between daily meal, plasma corticosterone and reward sensitization. Brain Res 695, 186–194 (1995).

49. Schmidt-Nielsen, B. & O’Dell, R. Structure and concentrating mechanism in the mammalian kidney. Am J Physiology-legacy Content 200, 1119–1124 (1961).

50. Trinh–Trang–Tan, M., Bouby, N., Kriz, W. & Bankir, L. Functional adaptation of thick ascending limb and internephron heterogeneity to urine concentration. Kidney Int 31, 549–555 (1987).

51. Sperber, I. Studies on the Mammalian Kidney. (1944).

52. Heisinger, J. F. & Breitenbach, R. P. Renal Structural Characteristics as Indexes of Renal Adaptation for Water Conservation in the Genus Sylvilagus. Physiol Zool 42, 160–172 (1969).

53. Brownfield, M. S. & Wunder, B. A. Relative medullary area: A new structural index for estimating urinary concentrating capacity of mammals. Comp Biochem Physiology Part Physiology 55, 69–75 (1976).

54. Gallardo, P. A., Cortés, A. & Bozinovic, F. Phenotypic Flexibility at the Molecular and Organismal Level Allows Desert-Dwelling Rodents to Cope with Seasonal Water Availability. Physiol Biochem Zool 78, 145–152 (2005).

55. Houtmeyers, A., Duchateau, L., Grünewald, B. & Hermans, K. Reference intervals for biochemical blood variables, packed cell volume, and body temperature in pet rats (Rattus norvegicus) using point-of-care testing. Vet Clin Path 45, 669–679 (2016).

56. Fitzsimons, J. T. The effects of slow infusions of hypertonic solutions on drinking and drinking thresholds in rats. J Physiology 167, 344–354 (1963).

57. Belcher, E. H. & Harriss, E. B. Studies of plasma volume, red cell volume and total blood volume in young growing rats. The Journal of Physiology 139, 64 78 (1957).

58. Busse, L. et al. The Detection of Visual Contrast in the Behaving Mouse. J Neurosci 31, 11351 11361 (2011).

59. Carandini, M. & Churchland, A. K. Probing perceptual decisions in rodents. Nat Neurosci 16, nn.3410 (2013).

60. Histed, M. H., Carvalho, L. A. & Maunsell, J. H. R. Psychophysical measurement of contrast sensitivity in the behaving mouse. J Neurophysiol 107, 758–765 (2012).

61. Haemisch, A., Guerra, G. & Furkert, J. Adaptation of corticosterone-but not β-endorphin-secretion to repeated blood sampling in rats. Lab Anim 33, 185–191 (1999).

62. Engelmann, M. et al. Endocrine and behavioral effects of airpuff-startle in rats. Psychoneuroendocrino 21, 391–400 (1996).

63. Plotsky, P. M. & Meaney, M. J. Early, postnatal experience alters hypothalamic corticotropin-releasing factor (CRF) mRNA, median eminence CRF content and stress-induced release in adult rats. Mol Brain Res 18, 195–200 (1993).

64. Akana, S. F. & Dallman, M. F. Chronic Cold in Adrenalectomized, Corticosterone (B)-Treated Rats: Facilitated Corticotropin Responses to Acute Restraint Emerge as B Increases. Endocrinology 138, 3249–3258 (1997).

65. Armario, A., Lopez-Calderon, A., Jolin, T. & Balasch, J. Response of anterior pituitary hormones to chronic stress. The specificity of adaptation. Neurosci Biobehav Rev 10, 245–250 (1986).

66. Kalinichev, M., Easterling, K. W., Plotsky, P. M. & Holtzman, S. G. Long-lasting changes in stress-induced corticosterone response and anxiety-like behaviors as a consequence of neonatal maternal separation in Long–Evans rats. Pharmacol Biochem Be 73, 131–140 (2002).

67. Toth, L. A. & Gardiner, T. W. Food and water restriction protocols: physiological and behavioral considerations. Contemporary topics in laboratory animal science / American Association for Laboratory Animal Science 39, 9 17 (2000).

68. Nicholaidis, S. & Rowland, N. Long-term self-intravenous “drinking” in the rat. J Comp Physiol Psych 87, 1 (1974).

